# Waiting for the Perfect Vaccine

**DOI:** 10.1101/2024.02.07.579403

**Authors:** Gergely Röst, Zhen Wang, Seyed M. Moghadas

## Abstract

Vaccination has proven to be the most effective public health measure in the fight against various infectious diseases. For emerging or re-emerging diseases, a highly efficacious vaccine may not be available at the start of an outbreak. Timelines for availability of a safe and effective vaccine may significantly affect disease dynamics, its burden, and the healthcare resource utilization. Mitigating this impact may then rely on low-efficacy vaccines that may be rapidly produced and distributed to at-risk populations at the early stages of an outbreak. With the expectation for arrival of a more effective vaccine at a later stage of the outbreak, the optimal vaccination coverage with the existing, low-efficacy vaccines is elusive. While flattening the outbreak if a significant proportion of the susceptible population is vaccinated with a low-efficacy vaccine, the overall infections may not be minimized if a small proportion of the population left unvaccinated when a highly efficacious vaccine becomes available. The optimal coverage for early vaccination could thus depend on several parameters including the efficacy of the currently available vaccines, arrival timing of a more effective vaccine and its efficacy, and the transmissbility of the disease. Here, we develop a deterministic system of differential equations to investigate the optimal vaccination coverage with a low-efficacy vaccine within the aforementioned parameter space. Despite simplifying assumptions, we illustrate that minimizing the overall infections does not necessarily correspond to the highest coverage of early vaccination. However, a high vaccination coverage, even with a low-efficacy vaccine, may still contribute to alleviating severe disease outcomes and reducing healthcare resource utilization.

## 1 Introduction

Emerging infectious diseases continue to pose significant threats to the health and socioeconomic well-being of human populations [1, 2]. At the time of emergence, strategies to quell the spread of the invading pathogen often rely on traditional and non-pharmaceutical measures, such as those implemented during the early stages of the COVID-19 pandemic [3–8]. These interventions, if not effective to contain the disease at the source, may delay widespread infections until more effective and/or preventive measures such as therapeutics and vaccines become available [6, 9, 10].

Vaccines are an important pharmaceutical measure [11] that may become available after the characteristics of the emerging pathogen are identified. However, the effectiveness of vaccines may depend on various factors including the biology and epidemiology of the disease, attributes of the target population for vaccination, as well as technologies for vaccine development [12–14]. As experienced during the COVID-19 pandemic, it is possible to have several vaccines developed using different technologies [15–18], rendering different effectiveness against infection and disease outcomes [19–23]. Importantly, the timelines for availability of these vaccines could vary during an outbreak, depending on the vaccine technology used and manufacturing capacity.

Although target product profiles [24] often indicate the range and lowest point estimates for the efficacy of a vaccine to be recommended for mass immunization, even low efficacy vaccines with adequate safety profiles can still contribute to reducing the disease burden [25]. The possibility that a vaccine with low efficacy becomes available at the early stages of an emerging disease raises a number of questions for vaccination policies. For example, it is unclear what proportion of the population should be vaccinated to minimize infections if a more effective vaccine is expected to arrive at a later date. On one hand, increasing the vaccination coverage with the current vaccine reduces the risk of disease transmission in the population and decelerates the spread of disease until a more effective vaccine becomes available. On the other hand, it reduces the proportion of the population unvaccinated and eligible to receive the second, more effective vaccine. The optimal coverage for early vaccination could thus depend on several parameters including the efficacy of the currently available vaccines, transmissbility of the disease, and the arrival timing of a more effective vaccine and its efficacy. When timelines for availability of the second vaccine is unknown, the efficacy of the current vaccine may play a significant role in determining the optimal proportion of the population that should be vaccinated to minimize the overall infections. Our study here aims to investigate this optimal scenario using a simple deterministic transmission dynamic model with simplifying assumptions.

## 2. Methods

We consider a deterministic system of differential equations, where a population of size *N* is divided into the four compartments of susceptible, vaccinated, infected (and infectious), and recovered individuals. We assume that a proportion *p* of the population is vaccinated at the onset of the outbreak, with a vaccine (*V*_1_) that provides an efficacy of *ϵ*_1_ < 1 against infection. We also assume that a second vaccine (*V*_2_) becomes available at some time *T* > 0 during the outbreak. For simplicity, we assume that *V*_2_ is a perfect vaccine preventing infection in all vaccinated individuals. At time *T*, we consider susceptible individuals who have not received *V*_1_ to be vaccinated with *V*_2_, thereby preventing any new infections among these individuals. However, those who have been vaccinated with *V*_1_ may still become infected at *t* > *T* due to imperfect efficacy of *V*_1_. With these assumptions, the dynamics of disease spread on the time interval [0, *T*] can be expressed by

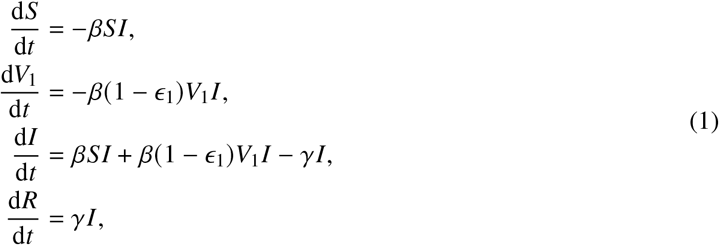

where *β* is the rate of disease transmission per contact per time, and *γ* is the recovery rate of infected individuals. For *t* > *T*, the model reduces to the following system

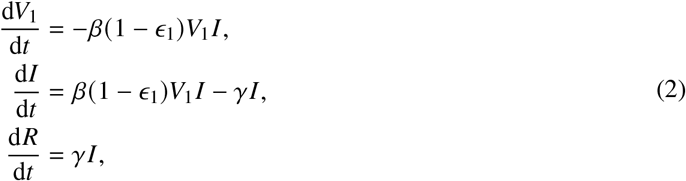

In the model (1)-(2), both recovered individuals and those vaccinated with *V*_2_ have a long-lasting immunity. At the onset of the outbreak, *V*_1_ (0) = *pN* and *S* (0) = (1 −*p*) *N*. Assuming an infected individual triggers the outbreak, simple calculation provides the expression for the reproduction number, given by

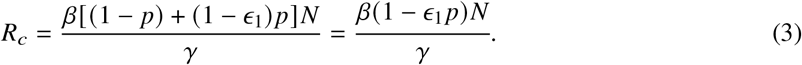

In the absence of any intervention, system (1) reduces to the classical SIR model with the reproduction number

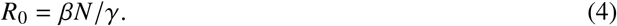

Thus, the control reproduction number with vaccination is *R*_*c*_ = (1 −*ϵ*_1_ *p*) *R*_0_. Based on the theory of mathematical epidemiology, if *R*_0_ < 1, the outbreak is expected not to take off, even without any interventions. Similarly, vaccination that achieves *R*_*c*_ < 1 prevents the outbreak. Our aim here is to determine the optimal *p* at which the total number of infections throughout the outbreak is minimized when *R*_0_ > 1 for given *ϵ*_1_ and *T* . There are two extreme cases for which the optimal *p* can be easily determined. The first case corresponds to the situation where *T* = 0. In this case, given that *V*_2_ is a perfect vaccine, all susceptible individuals should be vaccinated with *V*_2_, rendering *p* = 0 as optimal proportion of the population that is vaccinated with *V*_1_. The second case relates to the scenario in which *T* is large compared to the timelines of the outbreak, essentially making *V*_2_ unavailable during the outbreak. In this case, the minimum number of infections occurs when all susceptibles are vaccinated with *V*_1_, i.e., *p* = 1 is optimal. However, if *T* is a time during the outbreak, the optimal *p* is unknown and would depend on both *T* and *ϵ*_1_. Here, we first explore this optimal *p* for some special cases through analytical approaches. Then, we determine the optimal *p* for the general cases through means of simulations. The model parameters, and their values used in later simulations are described in Table 1.

**Table 1.** Parameters and values applied in the model simulations. The values 2.4 × 10^−6^, 4 × 10^−6^, 6 × 10^−6^ and 8 × 10^−6^ of the transmission rate correspond to basic reproduction numbers 1.2, 2, 3 and 4, respectively.

| Parameter | Interpretation | Values |
| --- | --- | --- |
| $\beta$ | Transmission rate | $2.4 \times 10^{-6}$<br>$4 \times 10^{-6}$<br>$6 \times 10^{-6}$<br>$8 \times 10^{-6}$ |
| $\gamma$ | Recovery rate ($\text{day}^{-1}$) | 0.2 |
| $\epsilon_1$ | Efficacy of the first vaccine | $[0, 1]$ |
| $p$ | Proportion of the population vaccinated with the first vaccine | $[0, 1]$ |
| $T$ | Time at which the perfect vaccine becomes available (days) | 0 – 300 |
| $N$ | Total population | $10^5$ |

## 3. Results for some special cases

In this section we provide some analytic results for special cases of the parameters, determining the optimal coverage *p*. By optimal, we mean the value of *p* ϵ[0, 1] such that the total number of infected individuals throughout the course of the epidemic is minimized. It would be useful to find invariant quantities (i.e., first integrals) of system (1) and (2).

### Proposition 1

*If p* ϵ (0, 1), *then S*(*t*) > 0 *for t* ϵ (0, *T*) *and V*_1_(*t*) > 0 *for all t* > 0. *Moreover, V*_1_(∞) := lim_*t*→∞_ *V*_1_(*t*) > 0 *and* lim_*t*→∞_ *I* (*t*) = 0.

*Proof*. Integrating the first and second equations of (1), we obtain

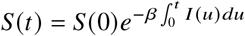

and

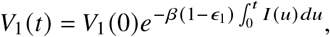

showing that they remain positive if their initial values are positive. Similarly,

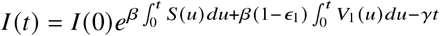

remains positive, and therefore, *S*(*t*) and *V*_1_(*t*) are monotone decreasing. Since *V*_1_(*t*) is non-negative, its limit exists. Since individuals can be infected only once in our model, it also holds that 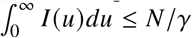 thus

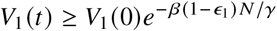

and *V*_1_(∞) > 0. Note also that for *t* > *T*,

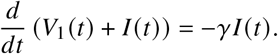

Since solutions are bounded between 0 and *N*, their derivatives are also bounded. If lim sup_*t*→∞_ *I* (*t*) > 0, then the integral of *I* (*t*) is unbounded (since its derivative is bounded), and then *V*_1_(*t*) + *I* (*t*) would become negative at some finite time which is not possible. Hence, lim_*t*→∞_ *I* (*t*) = 0.

### Proposition 2

*For S, V*_1_ > 0, *the following relations hold:*

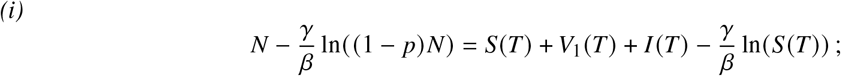

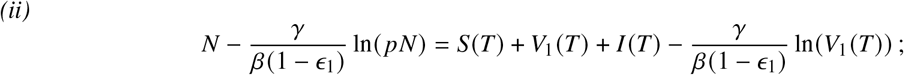

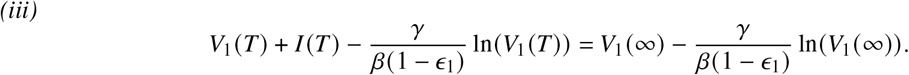

*Proof*. Define

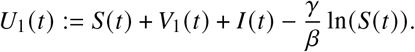

It is straightforward to check that 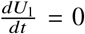, thus *U*_1_(*t*) is indeed invariant. Hence, *U*_1_(0) = *U*_1_(*T*), which proves *(i)*. Similarly, let

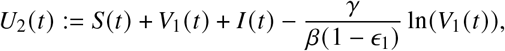

which is invariant, since 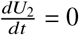. Hence,*U*_2_(0) = *U*_2_(*T*), that is (*ii*).

At *t* = *T*, all susceptibles are vaccinated by the perfect vaccine, hence *S* (*T*^+^) = 0, and for *t* > *T* the dynamics follow (2). For (2), one can check that

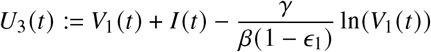

is also invariant. Consequently, *(iii)* holds.

*Remark 1* Rigorously speaking, the initial value *I* (0) > 0, thus either *S*(0) < (1 − *p*) *N* or *V*_1_(0) < *pN*. However, for many of our calculations *I* (0) = 0^+^ is negligible compared to *N*, so to simplify the presentation, *I* (0) is omitted from some calculations.

The total number of infected individuals resulting from infection in the susceptible class is *S*(0) − *S*(*T*) = (1 − *p*) *N* − *S*(*T*), while the total number of individuals being infected from the *V*_1_ compartment is *V*_1_(0) −*V*_1_(∞) = *pN* −*V*_1_(∞). Therefore, the overall number of infected individuals that we aim to minimize is

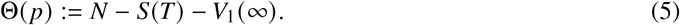

In this formula, both *S* (*T*) and *V*_1_ (∞) depend on *p*. The quantity Θ (*p*) / *N*, corresponding of the proportion of the population that will be infected throughout the epidemic, is also referred to as attack rate.

### 3.1 First vaccine with high efficacy

#### Proposition 3

*If ϵ*_1_ = 1 *then p* = 1 *is optimal*.

*Proof*. With *ϵ*_1_ = 1, 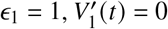, and (1) simplifies to

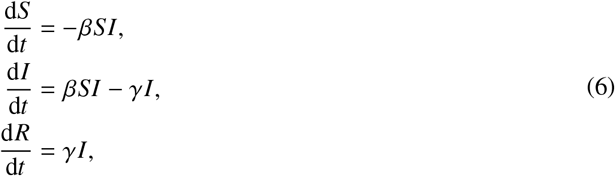

with *S*(0) = (1 − *p*) *N*, and *V*_1_(*t*) = *pN* for all *t*. In the case of *p* = 1, *S*(0) = 0 and *S*(*t*) = 0 for all *t*, in particular *S*(*T*) = 0. Since *ϵ*_1_ = 1, we always have *V*_1_(∞) = *V*_1_(0) = *pN* = *N*. Thus, Θ(1) = 0. On the other hand, if *p* < 1, then *S* is strictly decreasing and *S*(*T*) < *S*(0) = (1 − *p*) *N*, hence Θ(*p*) > 0.

The epidemiological interpretation of this result is that if the available vaccine at the onset of the outbreak is already perfect, then we would vaccinate everyone to prevent any infection. Any smaller coverage than 100% leaves some portion of the population susceptible to infection.

### 3.2 First vaccine with low efficacy

#### Proposition 4

*If ϵ*_1_ = 0 *then p* = 0 *is optimal*.

*Proof*. With *ϵ*_1_ = 0, system (1) becomes

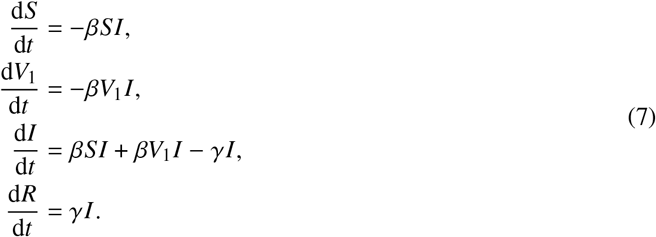

Now, we note that *S*(*t*) + *V*_1_(*t*) and *I* (*t*) are independent of *p*. In particular, *S*(*T*) + *V*_1_(*T*) and *I* (*T*) have the same value for all *p*. Moreover, for *p* = 0 we have *V*_1_(*T*) = 0, and for *p* > 0 we have *V* (*T*) > 0. The latter can be seen from *U*_2_(*t*) (see Proposition 2 *(ii*)), which gives

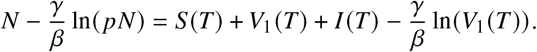

Rearranging yields

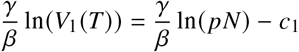

where *c*_1_ is a constant independent of *p*. This can be written as

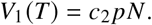

Thus, *V*_1_(*T*) − *V*_1_(∞) > 0, and

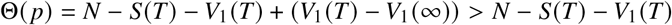

for *p* > 0; while for *p* = 0, we have *V* (*t*) = 0 for all *t*. Thus, Θ(0) = *N* − *S*(*T*) − *V*_1_(*T*), and we find *p* = 0 to □ be optimal.

The epidemiological interpretation of this result is that if the available vaccine offers very little protection, then we should not use it.

### 3.3 Short waiting time for the perfect vaccine

#### Proposition 5

*If T* = 0 *then p* = 0 *is optimal*.

*Proof*. In this case, all susceptibles are vaccinated immediately with the perfect vaccine, and the system (1) reduces to (2) with *V*_1_(0) = *pN* and *S*(0) = 0. If *p* > 0, then *V*_1_(∞) > 0 too, while for *p* = 0 we have □*V* (*t*) = 0 for all *t*. Since Θ(*p*) = *N* − *S*(*T*) − *V*_1_(∞) = *N* − *V*_1_(∞), we find that *p* = 0 is optimal.

The epidemiological interpretation of this result is that if the perfect vaccine becomes available very early, we should not use the weaker vaccine at all.

### 3.4 Large waiting time for the perfect vaccine

#### Proposition 6

*If T* = ∞ *then p* = 1 *is optimal*.

*Proof*. If *T* = ∞, then we have system (1) for all *t*, which is the same system considered in [26]. In the supplementary material of [26], the authors showed that the total number of infected individuals is a □ monotone decreasing function of *p*, hence *p* = 1 is optimal.

The epidemiological interpretation of this result is that if the perfect vaccine becomes available very late, we should vaccinate the whole population with the available weaker vaccine.

## 4. Results for the general case

Determining the optimal *p* in the general case can only be investigated by means of simulations. To do so, we chose a specific set of parameters by setting *β* = 4 × 10^−6^ and *γ* = 0.2, corresponding to a 5-day infectious period. For a population of *N* = 100, 000 individuals, and in the absence of vaccination, *R*_0_ = 2. We also varied the transmission rate to explore the variation in the optimal *p* when the disease exhibits lower (*R*_0_ = 1.2) or higher (*R*_0_ = 3, 4) transmissibility.

We simulated the model to determine the optimal *p* at which the attack rate (i.e., the proportion of the population infected throughout the outbreak) is minimized. As illustrated in Fig. 1, the optimal *p* varies as a function of *ϵ*_1_ and *T* . For sufficiently small *T*, the optimal *p* is a monotonic function and increases as *ϵ*_1_ increases. However, with longer delay in the availability of *V*_2_, the optimal *p* remains high for relatively low or relatively high *ϵ*_1_, but reduces for some intermediate levels of *ϵ*_1_. For example, at *T* = 100, the optimal *p* is 0.96 for both *ϵ*_1_ = 0.19 and *ϵ*_1_ = 0.48. This optimal *p* reduces to 0.84 when *ϵ* = 0.37. If *T* is sufficiently large, *p* = 1 is optimal, corresponding to vaccinating the entire susceptible population with *V*_1_.

**Fig. 1.**
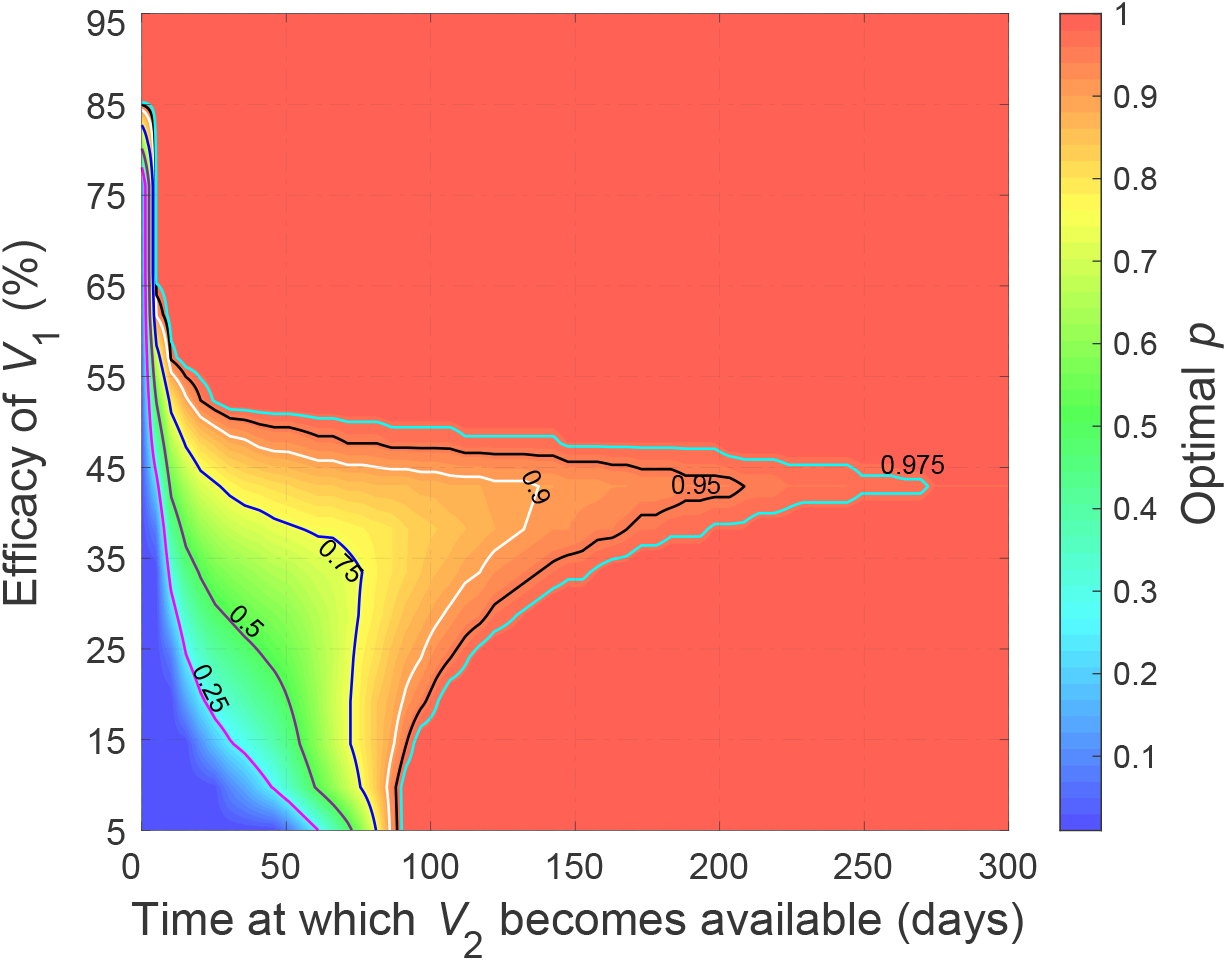
Optimal *p* for minimizing the total number of infections as a function of *ϵ*_1_ and *T*. The reproduction number is *R*_0_ = 2.

We also simulated the model to evaluate the effect of the reproduction number on optimal *p*. For a low reproduction number (*R*_0_ = 1.2), the optimal *p* presents a monotonic behaviour with respect to both *ϵ*_1_ and *T* (Fig. 2A). For any given *T*, the optimal *p* increases as *ϵ*_1_ increases. However, for a more contagious disease with reproduction number *R*_0_ = 4 (Fig 2B), we observed non-monotonic behaviour for optimal *p* similar to the case of *R*_0_ = 2 presented in Fig 1.

**Fig. 2.**
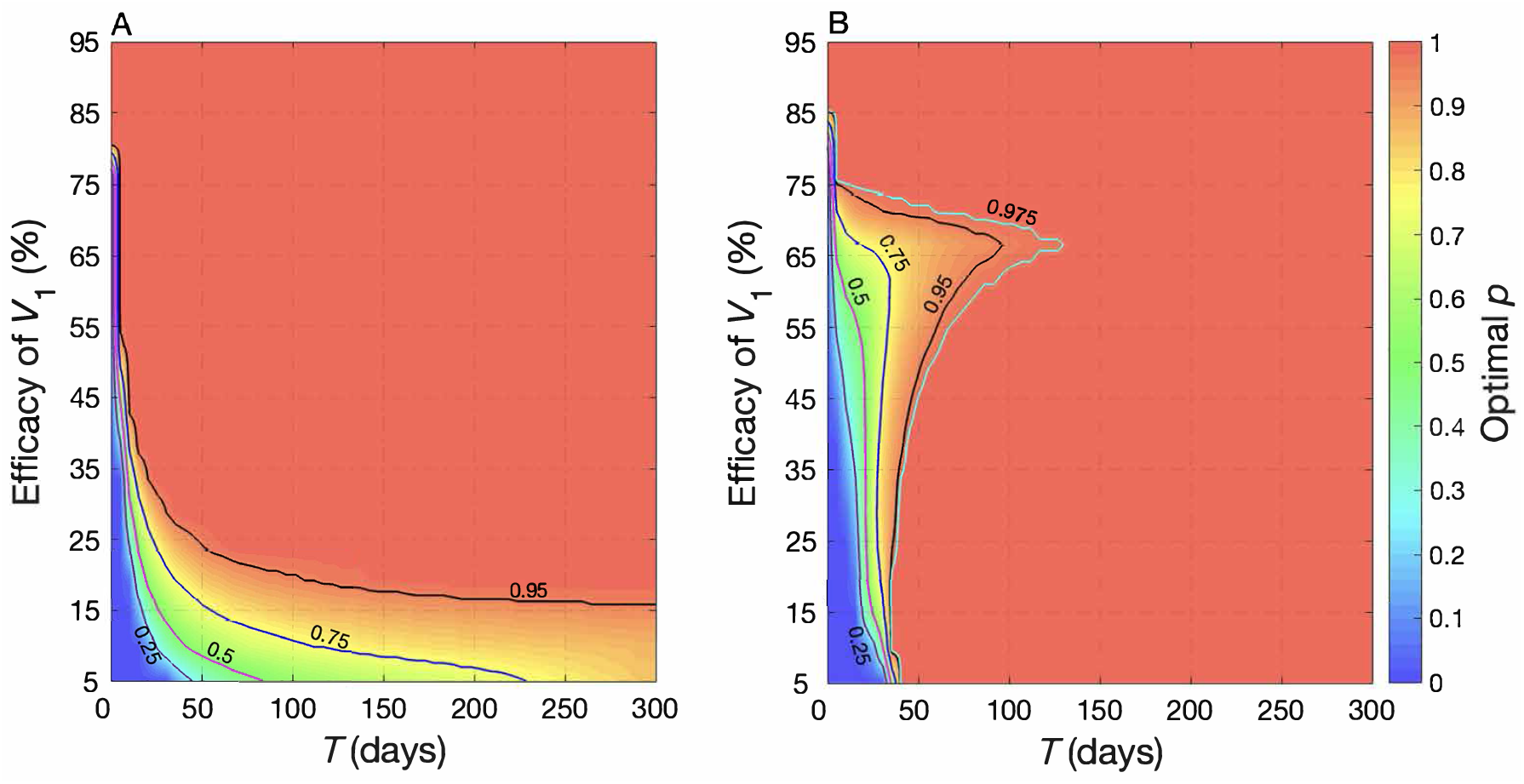
Optimal *p* for minimizing the total number of infections as a function of *ϵ*_1_ and *T* with (A) *R*_0_ = 1.2 and (B) *R*_0_ = 4 (B), corresponding to (A) *β* = 2.4 × 10^−6^ and (B) *β* = 8 × 10^−6^.

To illustrate the importance of optimal *p*, we simulated the incidence of infection when the efficacy of *V*_1_ in preventing infection was set to 0.25 (Fig 3A). In this case, the optimal *p* for reducing the overall attack rate is 0.556 assuming that *V*_2_ becomes available at *T* = 50 days after the onset of the outbreak. At this optimal *p*, the reproduction number derived from equation (3) is *R*_*c*_ = 1.722 at the onset of outbreak and the overall attack rate is 7.4% (Fig. 3B). We simulated the incidence with suboptimal *p* values of 0.5 and 0.6, which resulted in higher attack rates of 7.6% and 7.8%, respectively (Fig. 3B).

**Fig. 3.**
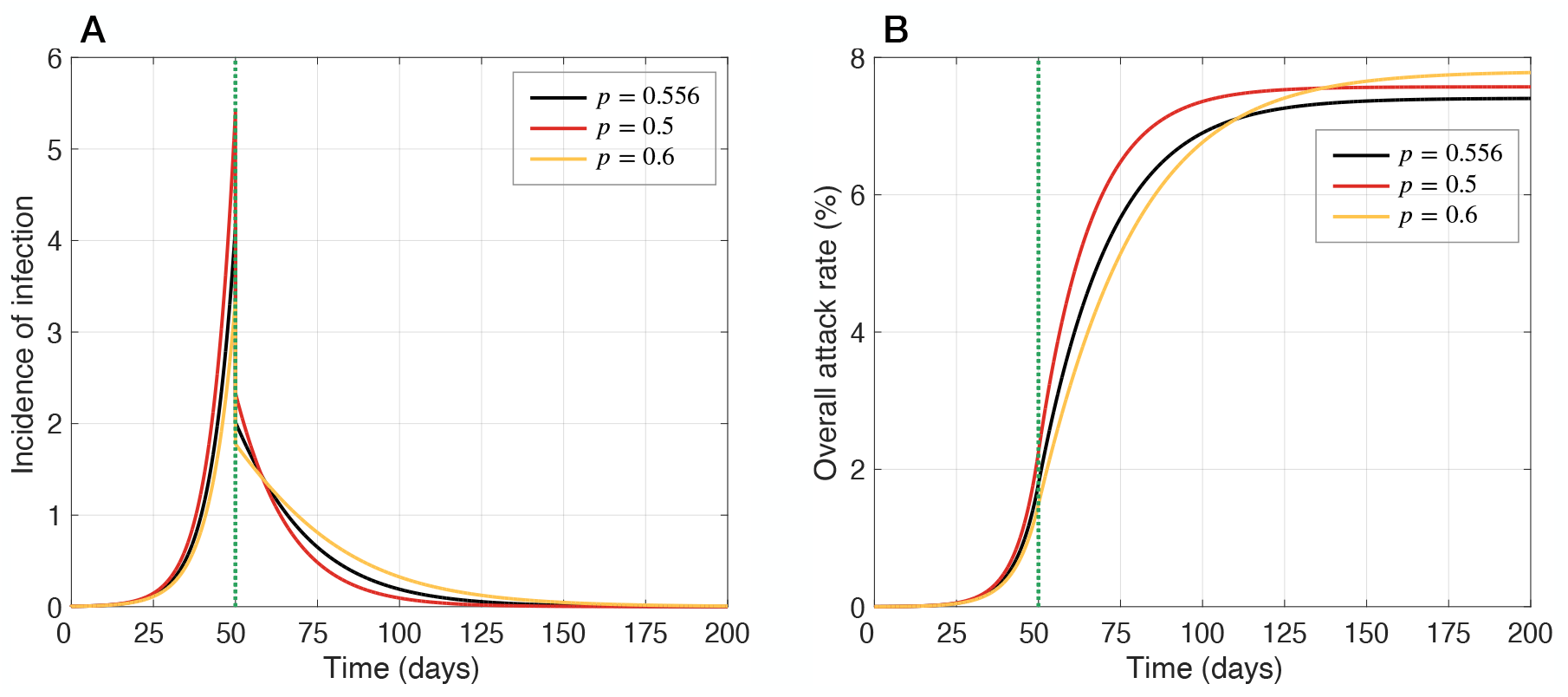
(A) Incidence of new infections at the optimal *p* = 0.556 (black curve), and suboptimal *p* = 0.5 (red curve) and *p* = 0.6 (orange curve). (B) Overall attack rate at the optimal *p* = 0.556 (black curve), and suboptimal *p* = 0.5 (red curve) and *p* = 0.6 (orange curve). Efficacy of *V*_1_ in preventing infection was assume to be *ϵ*_1_ = 0.25 and time for availability of *V*_2_ is *T* = 50 days after the start of the outbreak.

Clearly, as *p* increases, the spread of disease is decelerated, which reduces the attack rate for *t* < *T* . However, given that individuals vaccinated with *V*_1_ are not eligible to receive *V*_2_, the proportion of susceptible individuals who are not infected before *T* and would be eligible for *V*_2_ vaccination will also be reduced. This effectively translates to a higher attack rate for *t* > *T* due to a low efficacy of *V*_1_ (Fig 4). This non-linearity in optimal *p* illustrates the complexity of disease dynamics when multiple vaccines with different characteristics are expected to arrive during an outbreak.

**Fig. 4.**
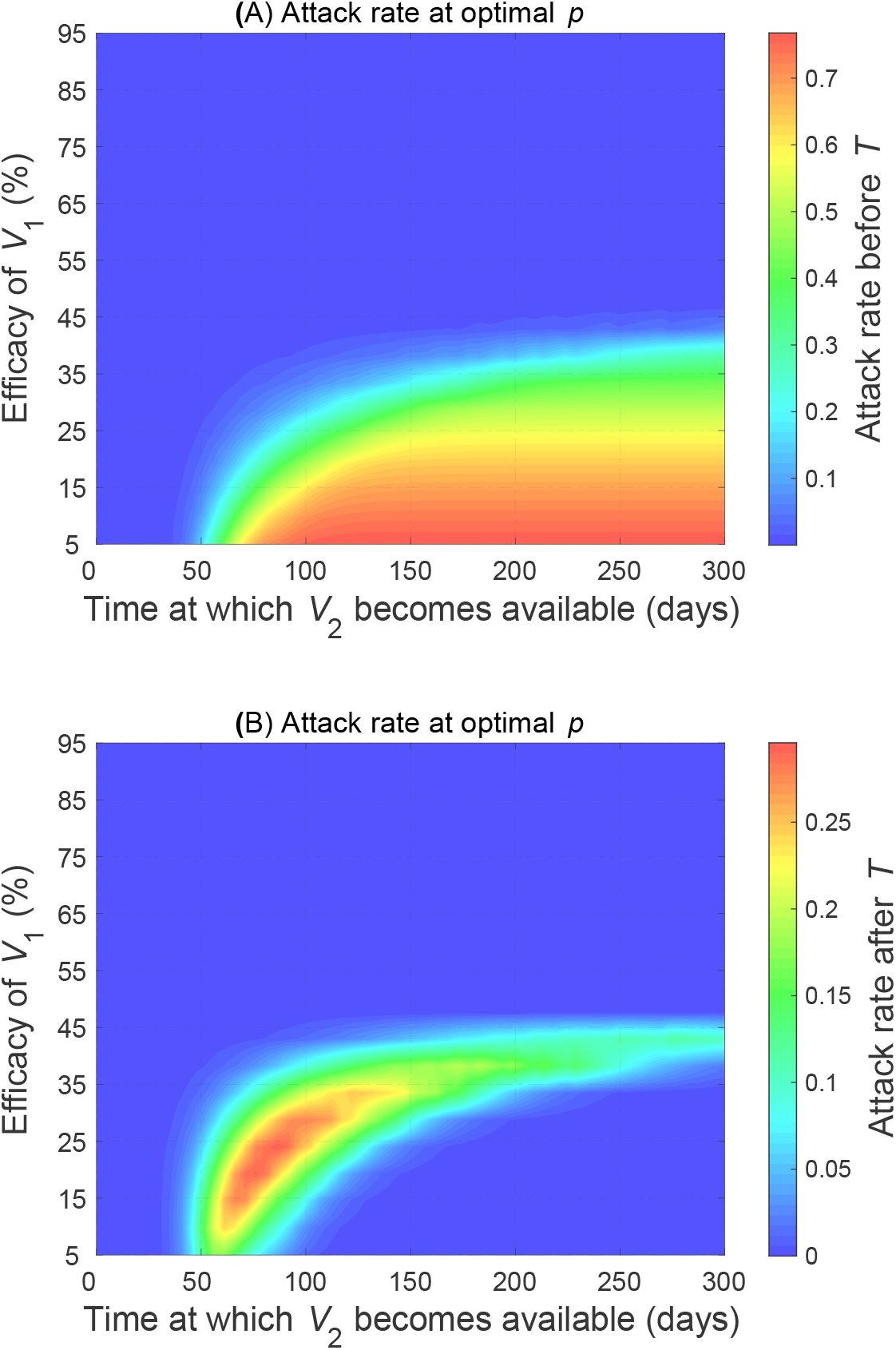
Attack rates as a function of *ϵ*_1_ and *T* before and after the availability of *V*_2_ at time *T* with the optimal *p* determined in Fig 1 for minimizing the total number of infections.

*Remark 2* From the expression for *R*_*c*_ = 1, it can be seen that at the threshold for disease control, the following relationship holds:

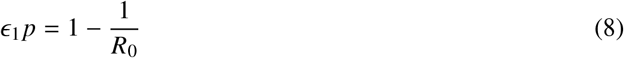

In the epidemiological theory for disease control, the expression (8) is referred to as herd immunity, which is defined as the proportion of the population needed to be immune against infection to prevent widespread. For example, when *R*_0_ = 2, if 50% of the susceptible population is protected, then a large outbreak is prevented. We compared the level of immunity at the onset of the outbreak derived from the optimal *p* (Fig 1) to this herd immunity threshold. Figure 5 shows that if the efficacy of *V*_1_ exceeds some threshold, the optimal *p* would raise the population-level protection above herd immunity level of 50% (corresponding to contour line of 0.5 in Fig 5), thus preventing a large outbreak.

**Fig. 5.**
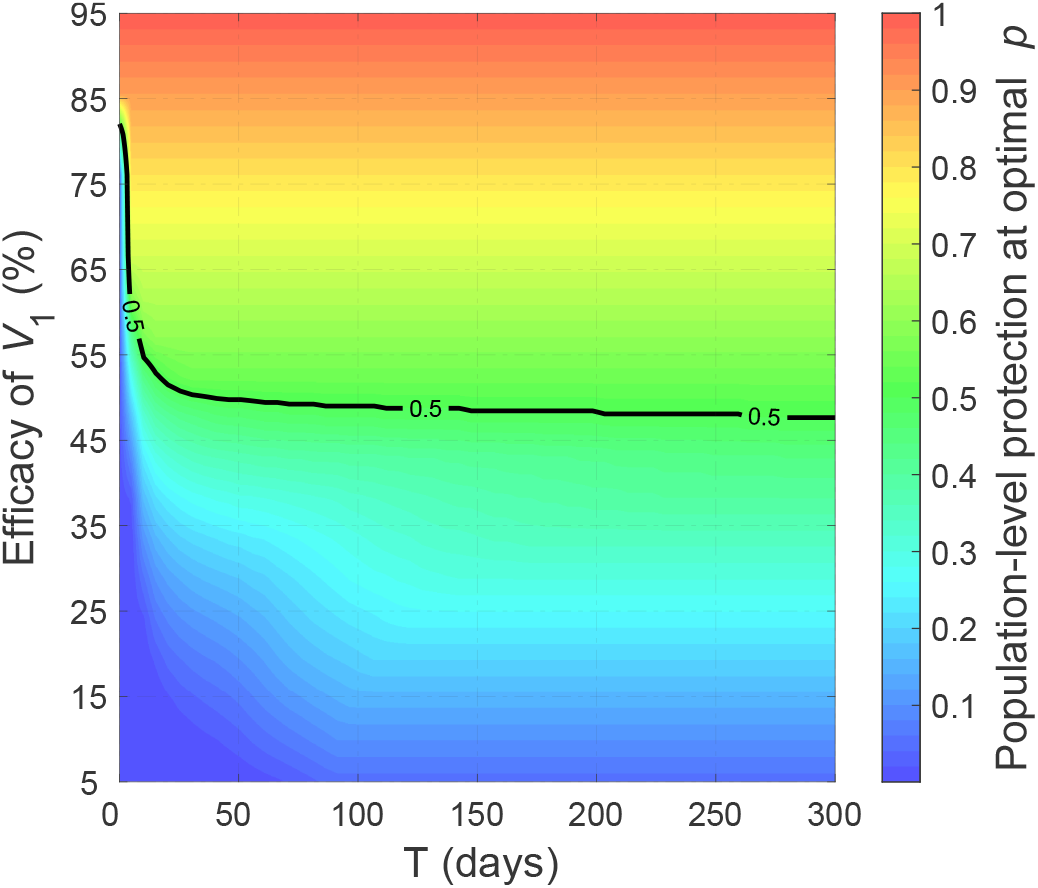
Population-level of immunity at the onset of the outbreak with the optimal *p* determined as a function of *ϵ*_1_ and *T* with *R*_0_ = 2.

We expanded our model to evaluate scenarios in which two additional vaccines become available during the outbreak at different times with different efficacy. Similar to the previous scenario, we assume that *V*_1_ is currently available with an efficacy of *ϵ*_1_ < 0.75. We consider the second vaccine (*V*_2_) to become available within the first 60 days of the outbreak, i.e., 0 < *T* ≤ 60, with a fixed efficacy of *ϵ*_2_ = 0.75 in preventing infection. We then assumed that the third vaccine (*V*_3_), with the simplifying assumption of being perfect in preventing infection, to be available on day 100 after the start of the outbreak. For this extension of the model, one may consider different proportion of the susceptible population being vaccinated with *V*_1_ and *V*_2_ to minimize the total number of infections. However, for the purpose of illustration, we simplify the parameterization and assume that the same proportion *p* of eligible individuals become vaccinated with *V*_1_ and *V*_2_ (when *V*_2_ becomes available). Unvaccinated individuals who are still susceptible on day 100 will then be vaccinated when *V*_3_ arrives.

Figure 6 shows that the optimal *p* depends not only on *ϵ*_1_ and *T*, but also on the reproduction number of the disease. When *R*_0_ is small (e.g., *R*_0_ = 1.2), the optimal *p* exhibits a monotonic behaviour and increases as *ϵ*_1_ or *T* increases. However, this pattern becomes irregular as *R*_0_ increases, rendering distinct regions of (*ϵ*_1_,*T*)-plane in which the same *p* is optimal with variation in other regions (Fig. 6).

**Fig. 6.**
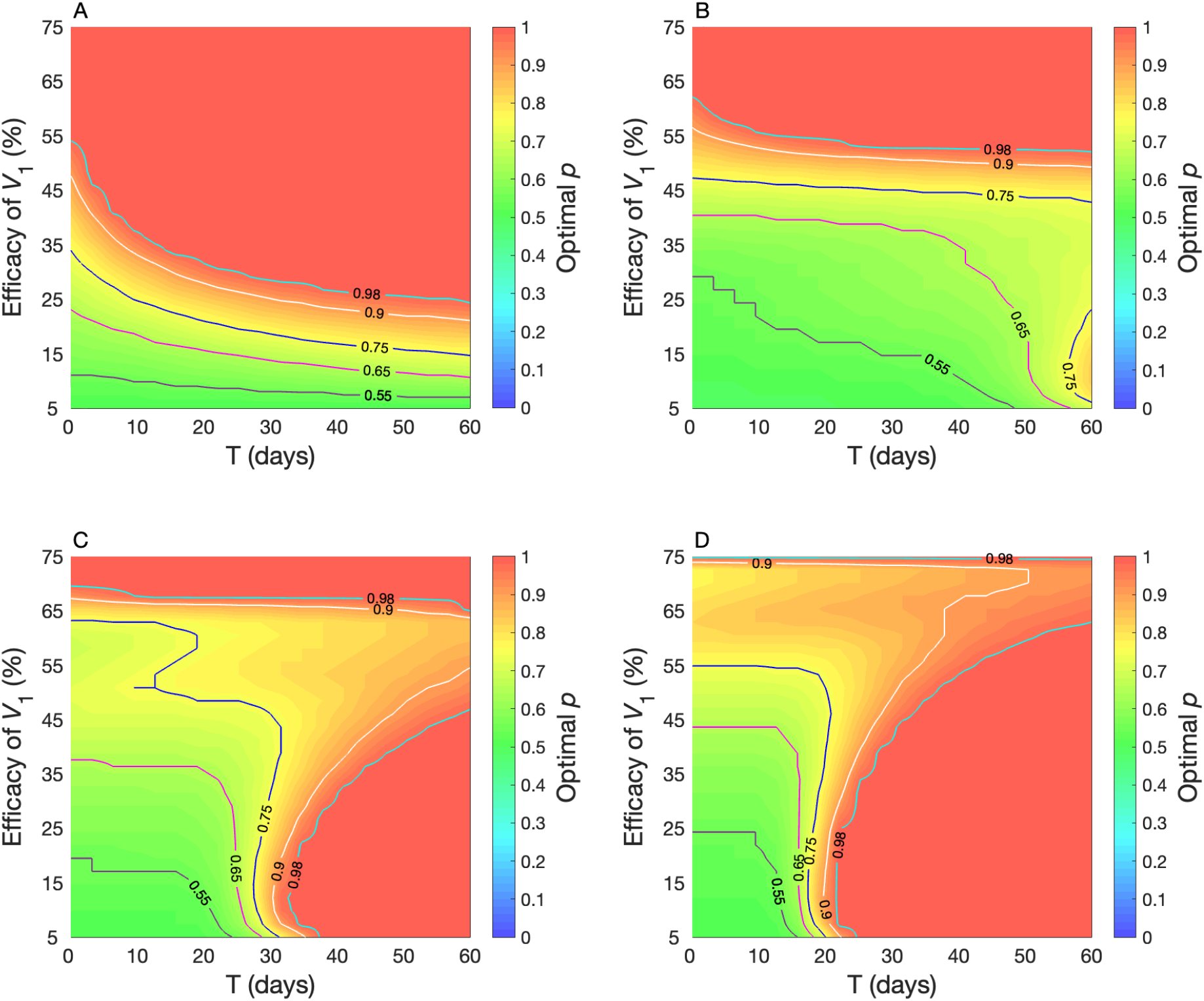
Optimal *p* for minimizing the total number of infections as a function of *ϵ*_1_ and *T*. The efficacy of second vaccine is fixed at 75% in preventing infection. The third vaccine with efficacy of 100% against infection is assumed to be available on day 100. Panels correspond to: *R*_0_ = 1.2 (A); *R*_0_ = 2 (B); *R*_0_ = 3 (C); *R*_0_ = 4 (D).

## 5 Discussion

In this study, we investigated optimal vaccination when multiple vaccines with different efficacies become available during an outbreak. Our results indicate that maximizing the coverage of vaccination at the onset of an outbreak does not necessarily lead to the lowest number of infections throughout the outbreak if more effective vaccines are expected to arrive at a later date. The optimality of early vaccination coverage could depend, as shown by our simulations, on the efficacy of vaccines, timelines for their availability and the reproduction number of disease indicating how fast the infection spreads through the population.

From a practical standpoint, our study may be more applicable to scenarios with the aim of minimizing the overall infections. However, the goals of public health extend beyond just minimizing the number of infections, and often include the reduction of severe disease outcomes (e.g., disease-related hospitalization and death) [25, 27–29]. In this context, early vaccination may be preferable to delayed vaccination using a product with higher efficacy [30]. More importantly, when healthcare resources are limited, the utilization of available capacity to reduce outcomes may be of critical importance to public health. Decelerating the spread of disease using available measures (such as vaccines) could alleviate the strain on the healthcare system, allowing for the available resources to be used for infected individuals at risk of severe outcomes. Flattening the outbreak with a maximum possible vaccination coverage early on in an outbreak, even if it leads to a higher attack rate over time, could still help mitigate healthcare burden and reduce potential deaths.

Our model uses various simplifying assumptions to reduce the complexity of disease dynamics. We assumed that a vaccine is available prior to the onset of an outbreak and a proportion of individuals can be vaccinated. This situation may not present itself in the case of an emerging disease. We also assumed that susceptible individuals can be vaccinated immediately when a vaccine becomes available. In practice, vaccination is a time-dependent process, and relies on the amount of vaccine supply and healthcare capacity to administer vaccines. We also did not consider the timelines needed for vaccination to induce immunity. Furthermore, we considered the availability of a vaccine with perfect efficacy during the outbreak. Although recent technologies (e.g., mRNA vaccines) provide promising platforms for highly efficacious vaccines to be developed in a relatively short period of time, no vaccine provides complete protection. The vaccine-induced immune protection is often affected by the individual attributes (e.g., health status and age), in addition to the characteristics of the pathogen.

Despite these limitations, our study provides an illustration of conundrum in complex vaccination dynamics. Future work is warranted to extend the model and investigate the optimality of vaccination coverage by relaxing the simplifying assumptions used in our model.

## Acknowledgements

GR was supported by Hungarian grants NKFIH KKP 129877, RRF-2.3.1-21-2022-00006, and TKP2021-NVA-09. SMM acknowledges the support from the Natural Sciences and Engineering Research Council of Canada, Discovery Grant; NSERC-MfPH Grant for Emerging Infectious Disease Modelling, and NSERC Alliance Grant.

